# Visual interference can help and hinder memory: Capturing representational detail using the Validated Circular Shape Space

**DOI:** 10.1101/535922

**Authors:** Aedan Y. Li, Keisuke Fukuda, Andy C. H. Lee, Morgan D. Barense

**Author notes:** Earlier versions of this work were presented at the Lake Ontario Visionary Establishment in 2016, the Canadian Society for Brain, Behaviour and Cognitive Science in 2017, the Vision Sciences Society in 2017 and 2018, the Toronto Area Memory Group in 2018, and the Memory Disorders Research Society in 2018. Portions of this work formed the Master’s thesis of AYL and a preprint is available at [placeholder]. Correspondence concerning this article should be addressed to Aedan Li, Department of Psychology, University of Toronto, 100 St. George Street, Toronto, ON, Canada, M5S 3G3. Contact.

## Abstract

Although we can all agree that interference induces forgetting, there is surprisingly little consensus regarding what *type* of interference most likely disrupts memory. We previously proposed that the similarity of interference differentially impacts the representational detail of color memory. Here, we extend this work by applying the *Validated Circular Shape Space* (Li et al., 2020) for the first time to a continuous retrieval task, in which we quantified both the visual similarity of distracting information as well as the representational detail of shape memory. We found that the representational detail of memory was systematically and differentially altered by the similarity of distracting information. Dissimilar distractors disrupted both fine- and coarse-grained information about the target, akin to memory erasure. In contrast, similar distractors disrupted fine-grained target information but increased reliance on coarse-grained information about the target, akin to memory blurring. Notably, these effects were consistent across two mixture models that each implemented a different scaling metric (either angular distance or perceived target similarity), as well as a parameter-free analysis that did not fit the mixture model. These findings suggest that similar distractors will help memory in cases where coarse-grained information is sufficient to identify the target. In other cases where precise fine-grained information is needed to identify the target, similar distractors will impair memory. As these effects have now been observed across both stimulus domains of shape and color, and were robust across multiple scaling metrics and methods of analyses, we suggest that these results provide a general set of principles governing how the nature of interference impacts forgetting.

---

Although we can all agree that distracting information disrupts memory, there is surprisingly little consensus regarding what *type* of interference most likely induces forgetting. In classic list-learning paradigms, memory failure is more likely to occur when distractors are similar to target items (Conrad, 1963; Conrad & Hull, 1964; Dyne, Humphreys, Bain, & Pike, 1990; Logie, Della, Wynn, & Baddeley, 2000; Wickelgren, 1965; Wickelgren, 1966; Wickens, Born, & Allen, 1963). Supporting these results, experiments in the long-term memory literature typically find greater memory and discrimination failures as the similarity between complex stimuli increases (e.g., Abe et al., 2011; Holden, Toner, Pirogovsky, Kirwan, & Gilbert, 2013; Kim & Yassa, 2013; Leal, Ferguson, Harrison, & Jagust, 2019; Ngo, Lin, Newcombe, & Olson, 2019; Pidgeon & Morcom, 2014; Reagh & Yassa, 2014; Sievers, Bird, & Renoult, 2019; Stark & Stark, 2017; Trelle, Henson, Green, & Simons, 2017; Yeung et al., 2013; Watson & Lee, 2013). However, other experiments have demonstrated the exact opposite pattern of results. Experiments in the short-term memory literature show that high distractor similarity benefits memory performance, a finding consistently replicated across stimulus types ranging from simple lines to complex faces (e.g., Dube & Sekuler, 2015; Dube, Zhou, Kahana, & Sekuler, 2014; Jiang, Lee, Asaad, & Remington, 2016; Kahana, Zhou, Geller, & Sekuler, 2007; Lin & Luck, 2009; Mate & Baques, 2009; Morey, 2018; Oberauer & Lin, 2017; Rademaker, Bloem, Weerd, & Sack, 2015; Rademaker, Chunharas, & Serences, 2019; Sanocki & Sulman, 2011; Sims, Jacobs, & Knill, 2012; Son, Oh, Kang, & Chong, 2020; Souza & Skora, 2017).

To reconcile this seemingly contradictory set of findings, distractor similarity has been proposed to differentially impact representational detail (Sun, Fidalgo, et al, 2017). Rather than conceptualizing memory as an all-or-nothing process whereby items are only ever remembered or forgotten, a more complete picture of performance may be provided by conceptualizing responses along a continuum of fine- and coarse-grained information. Behavioral, computational, and neuroimaging evidence reveals that memories not only vary along a continuum of representational detail (Bays, Catalao, & Husain, 2009; Berens, Richards, Horner, 2020; Greene & Naveh-Benjamin, 2020; Korkki, Richter, Jeyarathnarajah, & Simons, 2020; Ma, Husain, & Bays, 2014; Yonelinas, 2013; Zhang & Luck, 2008; 2009), but that there are also distinct neural signatures tracking the amount and types of detail present in memory (Brunec, Moscovitch, & Barense, 2018; Cooper & Ritchey, 2019; Nilakantan, Bridge, VanHaerents, & Voss, 2018; Oh, Kim, & Kang, 2019; Rademaker et al., 2019; Richter, Cooper, Bays, & Simons, 2016; Stevenson et al., 2018; Wais et al., 2017; Wais, Montgomery, Stark, & Gazzaley, 2018). These issues were empirically examined using a continuous retrieval task (*Figure 1a*; Wilken & Ma, 2004), whereby distractor similarity differentially impacted memory for the color of objects (Sun, Fidalgo, et al, 2017). Dissimilar distractors disrupted both fine- and coarse-grained information about the target, resulting in nearly the complete loss of the original memory trace. In contrast, similar distractors disrupted fine-grained information but increased the reliance on coarse-grained information about the target, resulting in blurring of the original memory trace.

**Figure 1.**
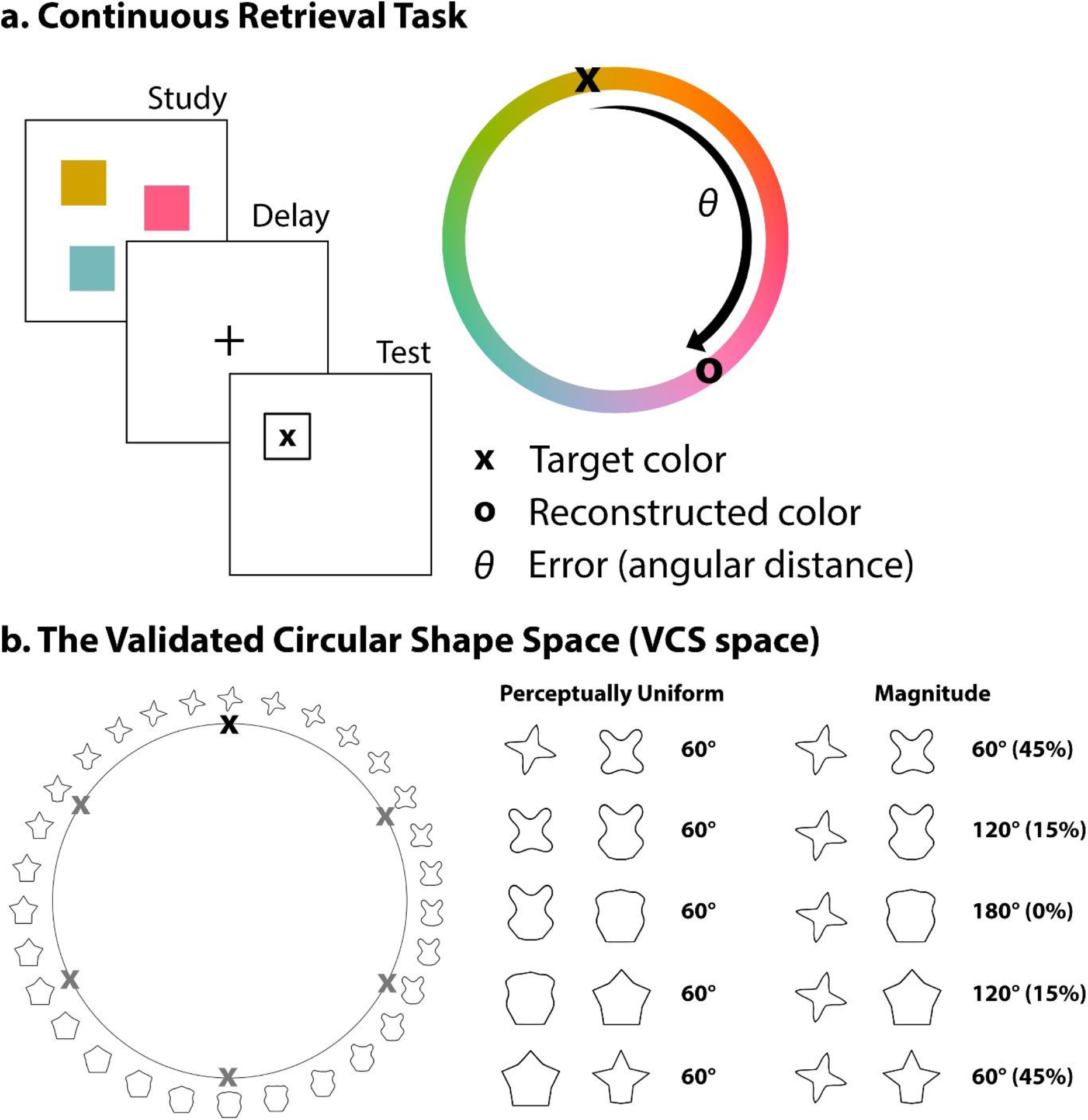
(*a*) Continuous retrieval task. During the study phase, target items are sampled from a random position on circular space. After a delay, participants reconstruct the target item on a 2D response circle during the test phase. Representational detail (i.e., error) is operationalized as the angular distance (θ) between a reconstructed and target item on circular space. (*b*) Visualization of Validated Circular Shape (VCS) space (note that only a subset of the 360 shapes are displayed for illustrative clarity). There are two ways to consider visual similarity on VCS space: 1) it is *perceptually uniform*, meaning that two shapes sampled from any distance (60 degrees apart on the example above) are about as similar as any two other shapes sampled from the same distance, and 2) the *magnitude* of visual similarity can be approximated by angular distance on VCS space, meaning that pairs of shapes sampled closer in angular distance tend to be more similar than pairs of shapes sampled from further distances (percent match in visual similarity displayed in brackets). Adapted from Li, Liang, Lee, and Barense (2020).

These differential effects of distractor similarity can reconcile the seemingly contradictory findings within the interference literature. For example, the similarity of interference may help or hinder memory depending on whether fine- or coarse-grained representations are needed to solve a particular task. In some cases, similar distractors may help memory when the preserved coarse-grained information can serve as a categorical reminder of the target item (Hanczakowski, Beaman, & Jones, 2017; Hardman, Vergauwe, & Ricker, 2017;

Jiang et al., 2016; Lin & Luck, 2009; Mate & Baques, 2009; Morey, 2018; Panichello, DePasquale, Pillow, & Buschman, 2019; Sanocki & Sulman, 2011). In other cases, this benefit from similar distractors may instead revert to a memory impairment when coarse-grained information is not sufficient to discriminate between similar distractor and target items (Barense et al., 2012; Cann, McRae, & Katz, 2011; Roediger & McDermott, 1995; Watson & Lee, 2013; Yeung et al., 2013).

Although the differential effects of interference on representational detail were robust for color (Sun, Fidalgo, et al., 2017), it is not yet known whether these findings generalize to more complex domains, such as shape. Indeed, extending previous color findings to a novel domain would provide strong evidence in support of a domain-general set of rules governing how interference impacts forgetting. For these reasons, we examined how distractor similarity impacted the representational detail of shape memory. In order to do so, however, we faced two nontrivial problems: 1) how could we manipulate the visual similarity of shape interference, and also 2) measure the representational detail of shape memory along a continuum of fine- and coarse-grained information?

## Quantifying Representational Detail Using a Circular Space

In order to study how interference impacts shape memory, we previously created the *Validated Circular Shape Space* (VCS space), a circular space whereby angular distance on a 2D circle is a proxy for visual similarity (Li, Liang, Lee, & Barense, 2020). VCS space was designed and validated to possess two properties (*Figure 1b*): 1) it is *perceptually uniform*, meaning that two shapes separated by any distance are about as visually similar as any two other shapes separated by the same distance, and 2) angular distance along the circle approximates the *magnitude* of visual similarity between two points, meaning that shapes sampled closer in angular distance are more visually similar than shapes sampled further apart in angular distance. For these reasons, VCS space is comparable to circular color spaces commonly used to quantify both visual color similarity (Lin & Luck, 2009; Sun, Fidalgo, et al., 2017) and the representational detail of color memory along a continuum (Bae et al., 2014; Brady, Konkle, & Alvarez, 2011; Fukuda, Awh, & Vogel, 2010; Hardman, Vergauwe, & Ricker, 2017; Ma, Husain, & Bays, 2014; Oberauer & Lin, 2017; Sun, Fidalgo, et al., 2017).

To measure the detail of representations, a circular space is then typically used in conjunction with a continuous retrieval task (*Figure 1a*). During the study phase of an experiment, a target item is displayed from a random position on a circular stimulus space. During the test phase, participants reconstruct the target item on a 2D response circle. An index of representational detail, *error*, is then calculated by taking the angular distance (θ) between the reconstructed and target item. Reconstructions low in error reflect a fine-grained memory for the target. That is, reconstructed items low in error are visually similar to the target item, and we can infer that these highly similar reconstructions contain abundant fine-grained information about the target. In contrast, reconstructed items higher in error are less visually similar to the target and therefore contain less fine-grained target information, instead consisting of primarily coarse-grained information about the target. Reconstructed items with the highest error bear little similarity to the target and subsequently contain very little fine- or coarse-grained information about the target.

Experimenters often fit mathematical models, such as *mixture models*, to data from a continuous retrieval task (*Figure 2a*; for more elaboration, see *Statistical Analysis – Fitting the mixture model*). The commonly used mixture model considers errors to originate from two sources (Zhang & Luck, 2008): random guesses, taken as evidence of retrieval failure, and the representational detail of the memory trace when the target is successfully retrieved. More specifically, the distribution of errors in angular distance are fit with two parameters that are thought to be independent (Ma et al., 2014; Suchow et al., 2013; Zhang & Luck, 2008): a uniform distribution (*g*) and a von Mises distribution (i.e., the circular analogue of a Gaussian distribution). The uniform distribution is thought to capture the probability of random guesses and is used to operationalize *retrieval success* (1 − *g*), because all errors are equally probable if observers do not remember any fine- or coarse-grained information about the target. In contrast, the von Mises distribution is thought to capture representational detail and is used to operationalize the *precision* of successfully retrieved memories (1/*sd*), whereby representations with more fine-grained information tend to be also lower in error across trials. Importantly, a particular advantage of the mixture model is the capability to dissociate the probability of retrieval success with the precision of the retrieved representation. For these reasons, the two-parameter mixture model has been incredibly influential (Zhang & Luck, 2008), informing debates regarding capacity limits in short-term memory (Fukuda, Awh, & Vogel, 2010; Fukuda, Vogel, Mayr, & Awh, 2010), forgetting over long timescales (Berens, Richards, & Horner, 2020), feature binding (Bays, Wu, & Husain, 2011; Golomb, L’Heureux, & Kanwisher, 2014), and dissociations between the number and quality of objects held in mind (Zhang & Luck, 2008; 2009).

**Figure 2.**
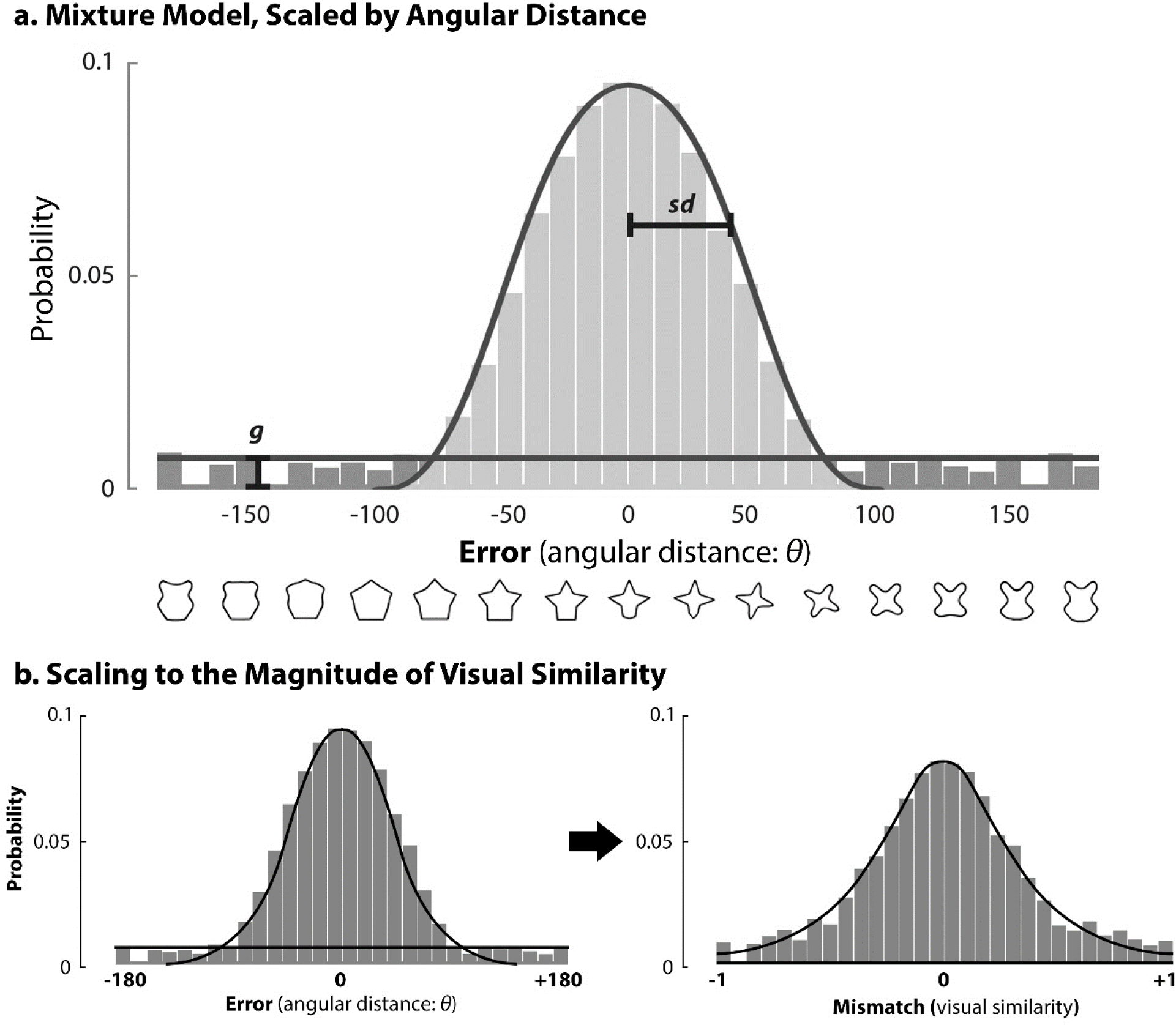
(*a*) The mixture model is commonly used to independently parameterize retrieval success with precision (Zhang & Luck, 2008). Error is first calculated as the signed angular distance between a target and a reconstructed item on circular space (*Figure 1a*). Zero error on the x-axis reflects an exact match between the target and reconstructed item, whereas the y-axis reflects the probability of reporting each error across all trials of the experiment. Two parameters are then fit to the distribution of errors using maximum likelihood estimation. The first parameter is a uniform distribution, thought to reflect the probability of random guesses (*g*). *P_mem_* is subsequently defined as retrieval success, 1 − *g*, representing the probability that some fine- or coarse-grained information about a target is successfully retrieved. In contrast, *precision* is defined as the inverse of the standard deviation (1/*sd*) of the second parameter, a von Mises distribution (i.e., the circular analogue of a Gaussian distribution), representing the representational detail of the successfully retrieved memory. Higher as opposed to lower precision reflects a memory containing more fine-grained information. (*b*) Scaling to the magnitude of visual similarity can influence the mixture model because the error distribution can change depending on the scaling metric. For these reasons, *P_mem_* and precision can be influenced by whether the data is scaled by angular distance or by visual similarity.

Despite the success of the mixture model, a separate body of work has conceptualized memory without the need to fit the uniform distribution used to derive the measure of retrieval success. This literature describes memory entirely as precision, such that responses with the highest error reflect memories with minimal detail rather than responses that necessarily reflect retrieval failure (Bays & Husain, 2008; Fougnie, Suchow, & Alvarez, 2012; van den Berg, Shin, Chou, George, & Ma, 2012; van den Berg, Awh, & Ma, 2014; Wilken & Ma, 2004). In other words, the responses with highest error which the mixture model would classify in terms of random guesses (i.e., as part of the uniform distribution *g*) are instead considered to be generated by representations with minimal detail.

This debate was reignited with the observation that angular distance on circular space *approximates* the magnitude of visual similarity but is not *identical* to visual similarity (Schurgin, Wixted, & Brady, 2019), meaning that angular distance does not reflect a one-to-one correspondence with the magnitude of visual similarity between two points on the circle. For example, although two items sampled from 60 degrees apart on the circular space are more visually similar than two items sampled from 120 degrees apart, the two items sampled from 60 degrees apart are not necessarily *twice* as similar as items sampled from 120 degrees (*Figure 1b*; Li et al., 2020). Importantly, this nonlinear relationship between the angular distance and the magnitude of visual similarity between two points on circular space may influence the interpretation of the mixture model, as the uniform distribution is fit to the data with the assumption that all errors on circular space are equally likely during a random guess (*Figure 2a*). Schurgin et al. (2019) showed that response distributions scaled by the magnitude of *perceived* visual similarity between the target and a participant’s response differed from response distributions that were scaled by the angular distance between the target and response (*Figure 2b*). More specifically, when responses were scaled in terms of their perceived visual similarity to the target, those responses with the highest error that had been classified as evidence of retrieval failure by the mixture model (i.e., those that appeared as part of the uniform distribution) were instead taken as evidence of responses with minimal memory strength (i.e., memories with minimal representational detail). That is, this new view holds that responses from a continuous retrieval task are best fit with a single parameter model that scales errors in terms of their perceived visual similarity to the target, rather than a two parameter mixture model that scales errors in terms of their angular distance from the target while independently characterizing retrieval success and precision. For these reasons, experimenters have suggested that the mixture model may be inaccurately parameterizing memory with separate retrieval success and precision parameters, as the errors used to fit the mixture model are not typically scaled to the magnitude of visual similarity (Bays, 2019; Schurgin et al., 2019).

## How Does the Type of Interference Influence the Detail of Representations?

In the present study, we used a continuous retrieval task in conjunction with mixture modelling to extend previous color findings into the domain of shape. To be consistent with our previous color work (Sun, Fidalgo et al., 2017), we used a mixture model to characterize retrieval success and precision. We directly accounted for the relationship between angular distance and the magnitude of visual similarity, comparing the results obtained from the mixture model to a parameter-free analysis that did not fit the mixture model. Using shapes whose visual similarity was explicitly quantified by VCS space (Li et al., 2020), we found that dissimilar interference disrupted both fine- and coarse-grained information, akin to memory erasure. In contrast, similar interference disrupted fine-grained but increased the reliance on coarse-grained information, akin to memory blurring. Critically, we replicate these findings across multiple methods of quantifying representational detail: 1) the mixture model fit to errors scaled by angular distance, 2) the mixture model fit to errors scaled by the magnitude of visual similarity, and 3) a set of parameter-free analyses that did not fit the mixture model. Across multiple converging methods of analyses, these results confirm that the similarity of distracting information differentially impact the representational detail of memory.

## Methods

### Participants

Thirty participants (*M_age_* = 20.13, *SD* = 2.16, *Females* = 21) were recruited from the undergraduate psychology student pool at the University of Toronto and from the community. This sample size was selected based on previous research (Sun, Fidalgo, et al., 2017), achieving 95% power to detect a main effect in a three-condition, one-way repeated measures analysis of variance (rm-ANOVA), assuming a medium-large effect size (*η*^2^ = 0.10) and a modest mean correlation between repeated measures (*r* = 0.40). All power analyses were conducted using G*Power (Faul et al., 2009). To increase power, we also increased the number of trials from 75 trials per interference condition (Sun, Fidalgo, et al., 2017) to 100 trials per interference condition (see *Procedure* section). Participants from the undergraduate psychology student pool were compensated with course credit and participants from the community were compensated with $20 CAD. This experiment required approximately 90 minutes to complete (*n* = 330 trials), which included consent, instructions, a short practice session of the task (12 trials), the actual experiment, and debriefing. Participants were allotted an optional self-timed break approximately every 15 minutes. This study was approved by the University of Toronto Ethics Board (protocol number 23778).

### Stimuli and Apparatus

The stimuli were shapes sampled from the *Validated Circular Shape Space* (VCS space, available on the Open Science Framework: https://osf.io/d9gyf/; Li et al., 2020), a circular shape space whereby angular distance is a proxy for visual similarity. The experiment code was developed in MATLAB using psychtoolbox-3 (Kleiner, Brainard, & Pelli, 2007). All stimuli were displayed on the monitor of a Latitude 3460 Dell laptop with a screen resolution of 1920 x 1080 and a frame refresh rate of 60 Hz. The monitor size was 12.18 inches by 6.85 inches, and the distance from the monitor to the participant was approximately 50 cm. All shape stimuli were presented on top of a uniform white background, subtending approximately 3.44 degrees of visual angle. Participants responded to on-screen instructions using the keyboard keys and the mouse.

### Procedure

We adapted a continuous task based on a previous experiment involving the color of objects (*Figure 3a*; Sun, Fidalgo, et al., 2017). At the beginning of every trial (*n* = 330), participants were shown a screen with the current trial number. When the space key was pressed by the participant, a central fixation cross appeared with “*Remember*” for 1000 ms. Each trial then contained three phases: study, interference (1-back task involving shapes of varying visual similarity to the target), and test, described in detail below.

**Figure 3.**
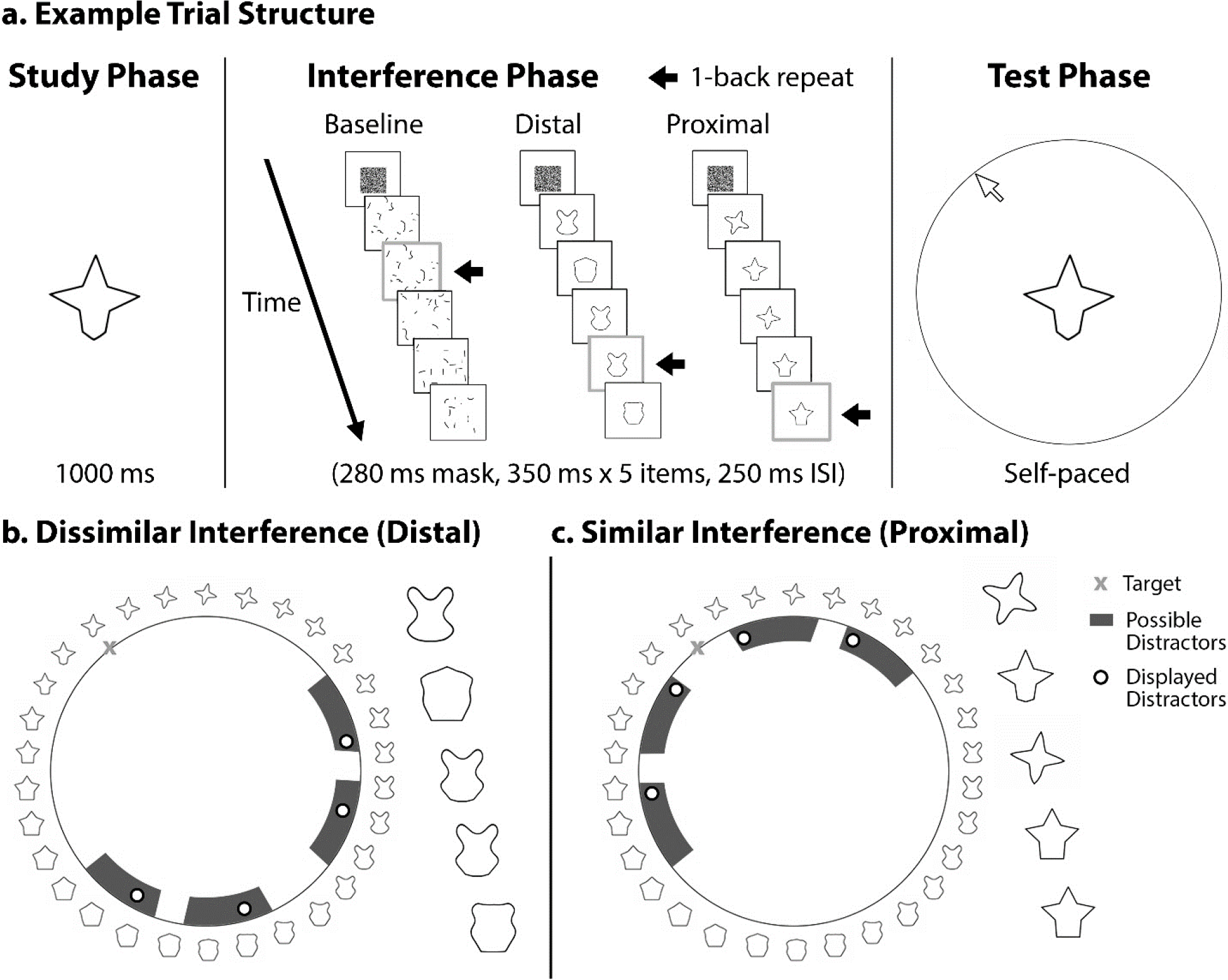
Example trial structure. (*a*) During the study phase, participants were asked to remember a target shape sampled from a random location on the Validated Circular Shape (VCS) space (Li, Liang, Lee, & Barense, 2020). Next, during the interference phase, distractors were presented sequentially in a 1-back task. There were three interference conditions: baseline (scrambled shapes), visually dissimilar (distal on circular space), and visually similar (proximal on circular space). During the test phase, participants reconstructed the target shape on a 2D response circle. Each degree of the response circle was associated with a shape from VCS space. This shape appeared at the center of the screen when the mouse cursor moved. (*b*) For trials in the Dissimilar Interference condition, distractors were sampled from the opposite side of VCS space relative to the target. (*c*) For trials in the Similar Interference condition, distractors were sampled from the same side of VCS space relative to the target. To control the variation in similarity, distractors were sampled from four cuts of circular space separated by 15 degrees distance. Distractors were never sampled from 14 degrees on either side of the target in the similar condition. Responses from this cut of circular space were used to determine the proportion of fine-grained responses in the parameter-free analysis.

Prior to the start of the experiment, participants were given detailed instructions and a set of 12 practice trials. Participants were explicitly asked: (1) to make their best guess if they could not remember the target shape, (2) to avoid verbally naming the shapes, (3) to be as accurate as possible, and (4) to press the space key upon identification of the repeating shape during the interference phase, which they were told could occur more than once during a given trial. In reality, only one repeat occurred during the n-back task. This instruction was provided to ensure that participants paid attention throughout the trial so that all interfering shapes were processed.

#### Study phase

Immediately following the “*Remember*” screen, a target shape was sampled randomly from the 360 shapes corresponding to each degree on VCS space, displayed for 1000 ms on the center of the screen.

#### Interference phase (1-back task)

After a 250 ms ISI, a grey-scale mask appeared for 280 ms to prevent visual afterimages. After another 250 ms ISI where nothing was presented on the screen, five distractor shapes (350 ms each, ISI 250 ms) were presented sequentially in three possible conditions: dissimilar (distractors distal on circular space), similar (distractors proximal on circular space), and baseline (scrambled shapes as distractors). As a measure of attention and to ensure that interfering material was processed, one of the distractor shapes repeated consecutively and participants were asked to identify this repeat (i.e., 1-back task). The temporal position of this repeat was randomly determined on every trial.

The type of interference varied across three conditions (*Figure 3b*), with the visual similarity of distractors explicitly quantified using the VCS space distance function (*Figure 9* in Li et al., 2020). The distance function relates degrees distance on VCS space with the exact magnitude of visual similarity (*Figure 1b*), with this function created from shape similarity judgments obtained from participants during the creation of VCS space. In the dissimilar condition, distractor shapes were sampled from a cut of 180 degrees on the opposite side of circular space relative to the target (i.e., 90 degrees to the left and right of a shape 180 degrees away from the target shape). Dissimilar distractors ranged from approximately 30% to 0% match in visual similarity with respect to the target. In the similar condition, distractor shapes were sampled from a cut of 180 degrees on the same side of circular space relative to the target (i.e., 90 degrees to the left and right of the target shape). Similar distractors ranged from approximately 85% to 30% match in visual similarity with respect to the target. In order to ensure that distractors were never identical to the target shape and that each distractor was visually distinct, the 180-degree slices in the dissimilar and similar conditions were further subdivided into four equal size subdivisions of 30 degrees, separated by “buffer zones” of 15 degrees distance from each other (buffer zones shown by the white segments in between the darker shaded regions in *Figure 3b*). The four subdivisions were each independently sampled in a random order to produce the overall set of distractor stimuli during each trial. By sampling distractors from these four subdivisions, we controlled the variation in visual similarity between distractors every trial, ensuring that interference was consistently similar in the similar condition and consistently dissimilar in the dissimilar condition.

Performance on the dissimilar and similar conditions was compared to a baseline “control” condition with distractor stimuli that were scrambled lines. Each of the dissimilar, similar, and baseline conditions contained 100 trials, and trials from each condition were presented in a randomized order.

Thirty trials with “random” interference were also interspersed to provide variability in the task and so that participants could not guess the trial type throughout the experiment. For these trials, distractor shapes were sampled from each quadrant of VCS space relative to the target item. These trials were not included in the statistical analysis because any potential impact of random interference could be due to a combination of effects caused by dissimilar and similar distractors.

#### Test phase

The test phase began with the mouse cursor centered on the screen, with participants instructed to reconstruct the target item along a 2D response circle. When the mouse cursor was moved by the participant, the specific shape stimulus that corresponded to the angle on VCS space occupied by the mouse cursor was displayed at the center of the screen (*Figure 3a*). All other shape stimuli were masked. The stimulus space was fully continuous, as the shapes were generated by morphing (see a visual example on the Open Science Framework: https://osf.io/msyw6/). Once a shape was reported through a mouse click, the next trial began.

Across participants, the shapes occupied different locations on the response circle during the test phase to ensure that there were no systematic relationships between particular shapes and locations. For example, if shape 1 was positioned on the response circle at 1 degree, shape 2 was positioned at 2 degrees, and shape 3 was positioned at 3 degrees for participant A, all the shapes were shifted by a random jitter for participant B (e.g. shape 1 may now be at 41 degrees, shape 2 at 42 degrees, shape 3 at 43 degrees, and so forth). These shape-location mappings were not jittered trial-by-trial because we reasoned that doing so would serve as an additional source of interference because the participant would have to extensively explore the space (and be exposed to uncontrolled interference) prior to being able to select a shape.

### Statistical Analysis

#### Fitting the mixture model

Error was first estimated by taking either the angular distance (θ) or the magnitude of visual similarity between a reconstructed and target item on circular shape space (*Figure 1*). We then fit the mixture model to errors across all trials within a condition using maximum likelihood estimation (*Figure 2*; Suchow et al., 2013), returning two parameters thought to be independent: a uniform distribution, *g*, and a von Mises distribution (i.e., the circular analogue of a Gaussian distribution). The uniform distribution *g* is thought to reflect the probability of random guesses (Zhang & Luck, 2008; 2009), because all errors on circular space are equally probable if observers do not remember any fine- or coarse-grained information about the target. To capture the probability that the memory is successfully retrieved, *P_mem_* is defined as 1 − *g*. In contrast, the von Mises distribution is thought to capture the representational detail of successfully retrieved memories by quantifying the spread of errors. Thus, the representational detail of the retrieved memory is defined as *precision*, modelled as the inverse standard deviation (1/*sd*) of the von Mises distribution. All model fitting analyses were conducted using MemToolbox (Suchow et al., 2013; the 2-parameter mixture model implemented using the *StandardMixtureModel* function).

We removed trials with incorrect or missing *n*-back responses to ensure that we included only trials in which we could verify that participants were paying attention to distractors (mean *n*-back = 94% in baseline, mean *n*-back = 91% in the dissimilar condition, and mean *n*-back = 91% in the similar condition). Next, the mixture model was fit to the data of every participant separately for the baseline, dissimilar, and similar conditions. For each mixture model analysis, two separate linear mixed models statistically analyzed *P_mem_* and precision as a function of interference condition (baseline, dissimilar, similar). To account for the within-subjects design, we modelled a random intercept for each participant (Pinheiro & Bates, 2000). The linear mixed models were estimated with an unstructured covariance matrix using the *lme* function from the *nlme* package (Pinheiro et al., 2018) in R version 3.6.1 (R Core Team, 2019), with statistical significance evaluated using the Satterthwaite approximation (Luke, 2017). Interference conditions were dummy coded in each linear mixed model (West, Aiken, & Krull, 1996), so that we compared the dissimilar to the baseline condition as well as the similar to the baseline condition. Effect sizes for individual comparisons were calculated using Cohen’s *d*, measured by the difference between condition means divided by the pooled standard deviation across conditions (Lakens, 2013).

#### Mixture model, scaled by angular distance

The mixture model was fit to errors scaled by angular distance, an analysis approach common in the literature (*Figure 3*; Ma et al., 2014; Zhang & Luck, 2008). Two measures were obtained from the mixture model: *P_mem_*, reflecting the probability of retrieval success, and *precision*, reflecting the representational detail of the memory trace when successfully retrieved.

#### Mixture model, scaled by the magnitude of visual similarity

Experimenters have suggested that responses classified as part of the uniform distribution (*g*) on the mixture model when scaled by their angular distance to the target may instead reflect a memory with minimal detail when scaled by their visual similarity to the target. As the mixture model analyses may be influenced by the scaling metric, we repeated the mixture model analyses after scaling responses in terms of their visual similarity to the target (*Figure 2b*).

We first rescaled errors using the VCS space distance function, a function converting each degree of distance on circular space directly into a percentage match in visual similarity between a participant’s response and the target item (*Figure 1b*). Errors in each interference condition were then replaced with the corresponding match in visual similarity to directly capture the magnitude of visual similarity between the reconstructed and target shapes. With this function, a 100% similarity score reflects maximal match in visual similarity, whereas 0% reflects minimal match in visual similarity with respect to VCS space. All scores were then subtracted from 100% to describe the resulting distribution in terms of visual mismatch, based on previous work on circular color space (Schurgin et al., 2019). We first accounted for the sign of the visual mismatch scores, then multiplied the distribution by 180 to describe the x-axis in the same dimensions as errors scaled by angular distance (*Figure 2*). Last, the mixture model was fit to these rescaled errors from each interference condition using MemToolbox (Suchow et al., 2013; the 2-component mixture model implemented using the *StandardMixtureModel* function).

#### Parameter-free analysis

In the final set of analyses, we examined performance without fitting the mixture model, instead interpreting response distributions directly along a representational continuum of fine- and coarse-grained information. This approach does not separate retrieval success with the precision of the memory trace like the mixture model, but instead considers representations along a single continuum of detail. Across all participants, errors were binned three ways within each interference condition: 1) fine-grained responses, 2) coarse-grained responses, and 3) responses with minimal or no detail (*Figure 6a*). Importantly, these three bins enabled us to investigate whether there were systematic differences in representational detail following exposure to different types of interference without fitting any model parameters, which could be influenced by the nonlinear relationship between angular distance and the magnitude of visual similarity. Using the VCS space distance function which relates angular distance to the magnitude of visual similarity (*Figure 1b*), we classified responses as fine-grained if they were made within 14 degrees on either side of the target (shapes in this region range from approximately 100% to 85% match in visual similarity to the target; Li et al., 2020). Note that these fine-grained reconstructions of the target reflect responses on VCS space where no distractors were ever sampled (see the white regions surrounding the target in *Figure 3b, c*). We classified responses as coarse-grained if they were made within 15 – 89 degrees on either side of the target, reflecting reconstructions that overlapped with the positions of distractors sampled from the similar condition (shapes in this region have an approximate 85% to 30% match in visual similarity to the target). We classified reconstructions as having minimal target information if they were made within 91 – 180 degrees on either side of the target, reflecting reconstructions that overlapped with the positions of distractors sampled from the dissimilar condition (shapes in this region have an approximate 30% to 0% match in visual similarity to the target). Responses made at 90 degrees from the target overlapped with the distractors sampled from both the dissimilar and similar conditions and were excluded (7 excluded trials out of a possible 7720 trials in the baseline, dissimilar, and similar conditions across all participants).

The proportion of responses made to each bin type was calculated separately for the baseline, dissimilar, and similar conditions, such that each participant had a mean value for each of the three bins for each of three interference conditions (i.e., 9 mean values in total for each participant). Collapsed across fine-grained, coarse-grained, and minimal target information bins, the proportion of responses made to each interference condition summed to 1. To conduct inferential statistics, we used a linear mixed model to examine the proportion of responses made as a function of bin type and interference condition. Note that there could be no main effect of interference condition at the omnibus level of the model, as the proportion of responses for each of the three interference conditions was always the same (i.e., 1). To account for the within-subjects design, a random intercept was modelled for each participant (Pinheiro & Bates, 2000), with statistical significance evaluated using the Satterthwaite approximation (Luke, 2017). Last, we tested a series of planned comparisons to examine the influence of distractor similarity on representational detail. Interference conditions were dummy coded (West, Aiken, & Krull, 1996) to compare the dissimilar to the baseline condition as well as the similar to the baseline condition for each bin type (fine-grained, coarse-grained, and minimal target information).

## Results

We examined how the type of distracting information impacted the detail of representations using separate mixture model analyses to characterize retrieval success, denoted by *P_mem_*, and the representation detail of the memory trace when it is successfully retrieved, denoted by precision. In addition, a final parameter-free analysis directly examined the representational detail of the memory trace without fitting the mixture model. Anonymized data files and commented analysis code is available on the Open Science Framework (https://osf.io/msyw6/).

### Mixture model, scaled by angular distance

Two participants performed at floor performance on the baseline condition and were subsequently excluded. Inclusion of outliers did not change the results.

The mixture model was fit to errors scaled by angular distance in each interference condition. A linear mixed model revealed that *P_mem_* (*Figure 4a, b*) was significantly influenced by interference condition (*F*_2,54_ = 19.20, *p* < 0.001, *partial R*^2^ = 0.42). Relative to the baseline (“control”) condition, *P_mem_* was significantly lower in the dissimilar condition (*t*_54_ = 5.52, *p* < 0.001, *d* = 0.54), but significantly higher in the similar condition (*t*_54_ = 2.82, *p* = 0.0067, *d* = 0.35). The second linear mixed model revealed that precision (*Figure 4c, d*) was also significantly influenced by interference condition (*F*_2,54_ = 13.42, *p* < 0.001, *partial R*^2^ = 0.33). Notably, however, the nature of the interference affected precision differently than it did *P_mem_*. Precision in the dissimilar condition was not significantly different compared to the baseline condition (*t*_54_ = 0.30, *p* = 0.77). In contrast, precision was significantly lower in the similar condition compared to the baseline condition (*t*_54_ = 4.23, *p* = 0.001, *d* = 0.54).

**Figure 4.**
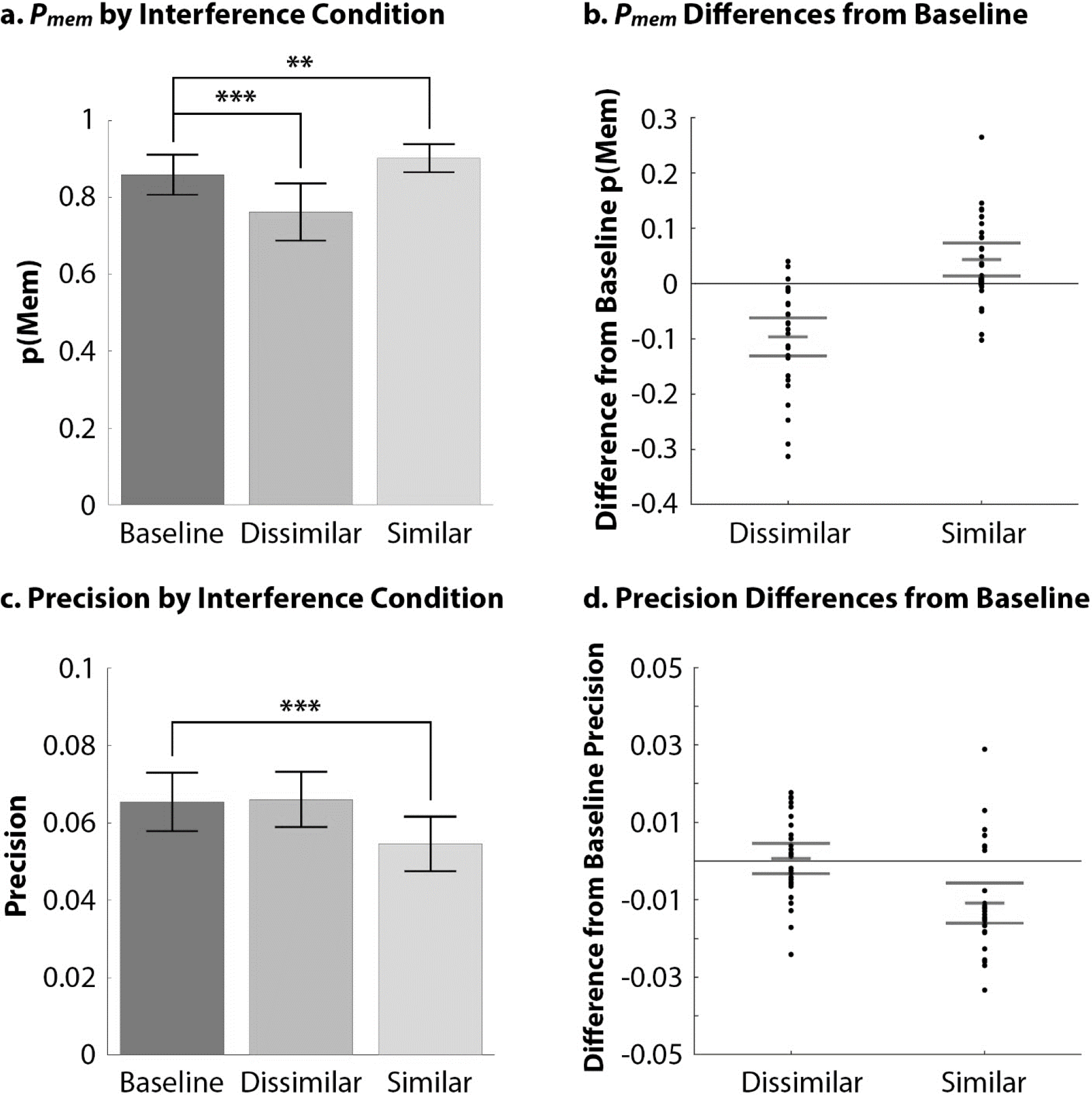
Mixture model fit to response distributions scaled by angular distance. (*a, b*) *P_mem_* was significantly reduced following visually dissimilar interference (distractors distal on circular space) relative to a baseline condition consisting of scrambled lines, and significantly increased following visually similar interference (distractors proximal on circular space) relative to the baseline condition. (*c, d*) In contrast, precision was not different between the dissimilar relative to the baseline condition, whereas precision was significantly lower in the similar relative to the baseline condition. Error bars in (*a, c*) reflect the 95% CI for the condition means. Error bars in (*b, d*) reflect the 95% CI for the within-subject differences between condition means from baseline. *** denotes *p* < 0.001, whereas ** denotes *p* < 0.01.

### Mixture model, scaled by the magnitude of visual similarity

To account for the fact that angular distance between two points on circular space does not reflect a one-to-one correspondence between their visual similarity, we repeated the mixture model analyses after scaling responses directly to their magnitude of visual similarity with the target (*Figure 1b*). The mixture model was then fit to each participant, obtaining a separate *P_mem_* and precision value for each interference condition (baseline, dissimilar, and similar). The response distribution for one participant after rescaling to the magnitude of visual similarity was found to be poorly fit with the mixture model (precision > 500000), so this participant was excluded.

Scaling the data by the magnitude of visual similarity revealed that *P_mem_* was significantly influenced by interference condition (*F*_2,52_ = 15.27, *p* < 0.001, *partial R*^2^ = 0.37) (*Figure 5a, b*). Relative to the baseline condition, *P_mem_* was significantly lower in the dissimilar condition (*t*_52_ = 5.00, *p* < 0.001, *d* = 0.46), but significantly higher in the similar condition (*t*_52_ = 2.34, *p* = 0.023, *d* = 0.22). The second linear mixed model revealed that precision (*Figure 5c, d*) was also significantly influenced by interference condition (*F_252_* = 10.01, *p* < 0.001, *partial R*^2^ = 0.28). Notably, however, the type of interference affected precision differently than it did *P_mem_*. Precision in the dissimilar condition was not significantly different compared to the baseline condition (*t*_52_ = 0.80, *p* = 0.43). In contrast, precision was significantly lower in the similar condition compared to the baseline condition (*t*_52_ = 4.10, *p* = 0.001, *d* = 0.74).

**Figure 5.**
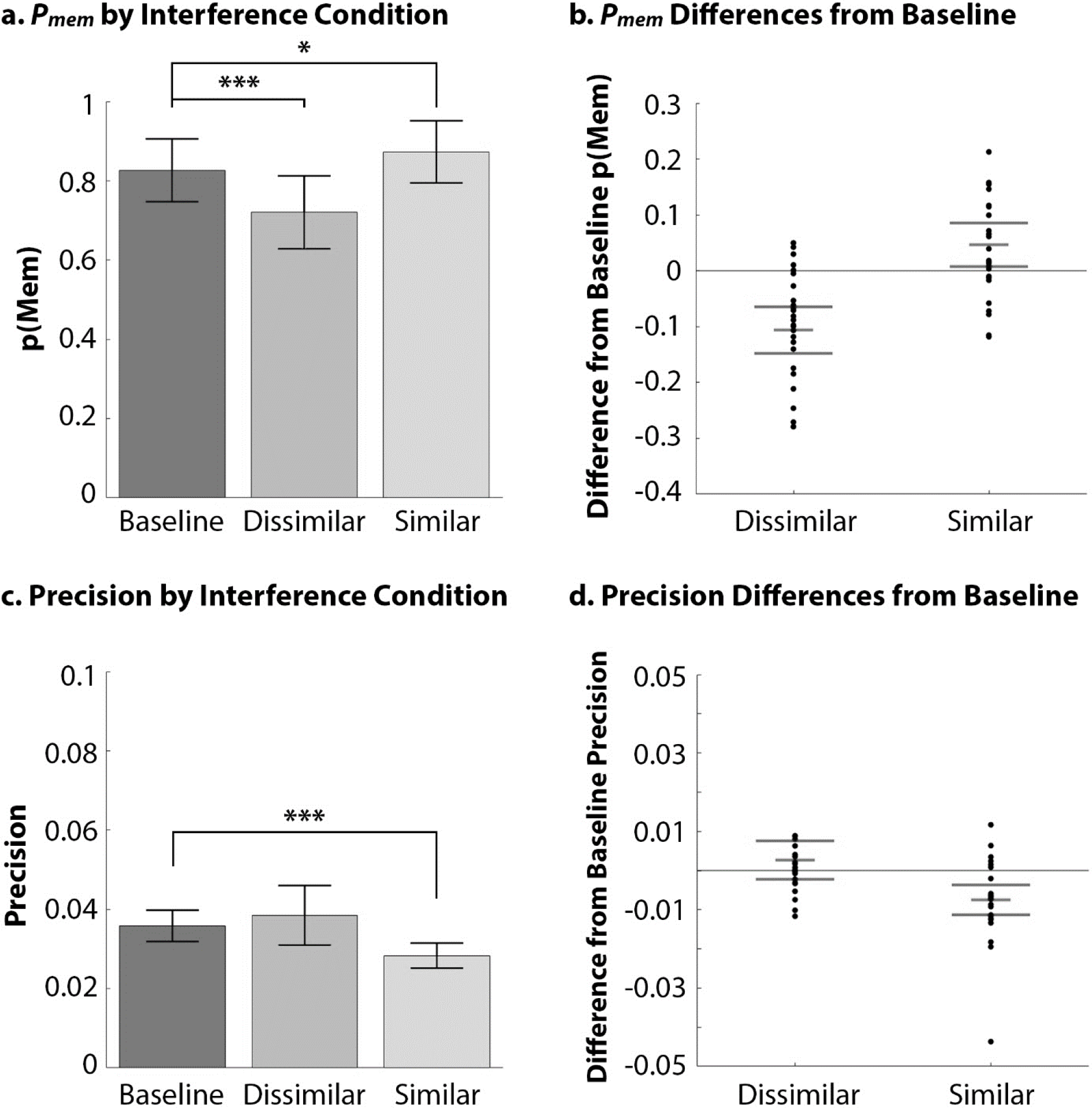
Mixture model fit to response distributions scaled by the magnitude of visual similarity. (*a, b*) *P_mem_* was significantly reduced following dissimilar interference (distractors distal on circular space) relative to a baseline condition consisting of scrambled lines, and significantly increased following similar interference (distractors proximal on circular space) relative to the baseline condition. (*c, d*) In contrast, precision was not different between the dissimilar relative to the baseline condition, whereas precision was significantly lower in the similar relative to the baseline condition. Error bars in (*a, c*) reflect the 95% between-subject CI for the condition means. Error bars in (*b, d*) reflect the 95% CI for the within-subject differences between condition means from baseline. *** denotes *p* < 0.001, whereas * denotes *p* < 0.05.

The mixture model fit to responses scaled by the magnitude of visual similarity (*Figure 5*) replicated results from when the mixture model was fit to responses scaled by angular distance (*Figure 4*). Not only does the type of interference differentially impact retrieval success and precision, we confirm that this finding was not driven by the nonlinear relationship between angular distance and the magnitude of visual similarity between two points on circular space (Schurgin et al., 2019).

### Parameter-free analysis

Last, we quantified representational detail without fitting model parameters (*Figure 6*). We binned responses in three ways for each condition based on the level of target-related detail the response contained: (1) fine-grained responses corresponding to approximately 100% to 85% match in visual similarity; (2) coarse-grained responses corresponding to approximately 85% to 30% match in visual similarity; and (3) responses with minimal or no detail, corresponding to approximately 30% to 0% match in visual similarity. Separate linear mixed models examined the influence of interference condition on each bin type. Note that we could not examine the main effect of interference condition collapsed across bin type, as the proportion of responses totaled to 1 within each condition (see *Experiment 1 – Statistical Analysis. Parameter-free analysis* section).

The linear mixed model found a significant main effect of bin type (*F*_2,216_ = 341.12, *p* < 0.001, *partial R*^2^ = 0.76), suggesting that collapsed across interference conditions, participants made responses at different proportions to the bins with fine-grained, coarse-grained, or minimal information. There was a significant interaction between bin type and interference condition, suggesting that the detail of representations was differentially influenced by the type of interference (*F*_4,216_ = 3.38, *p* = 0.011, *partial R*^2^ = 0.059). We conducted planned comparisons to directly examine the impact of interference condition on representational detail.

**Figure 6.**
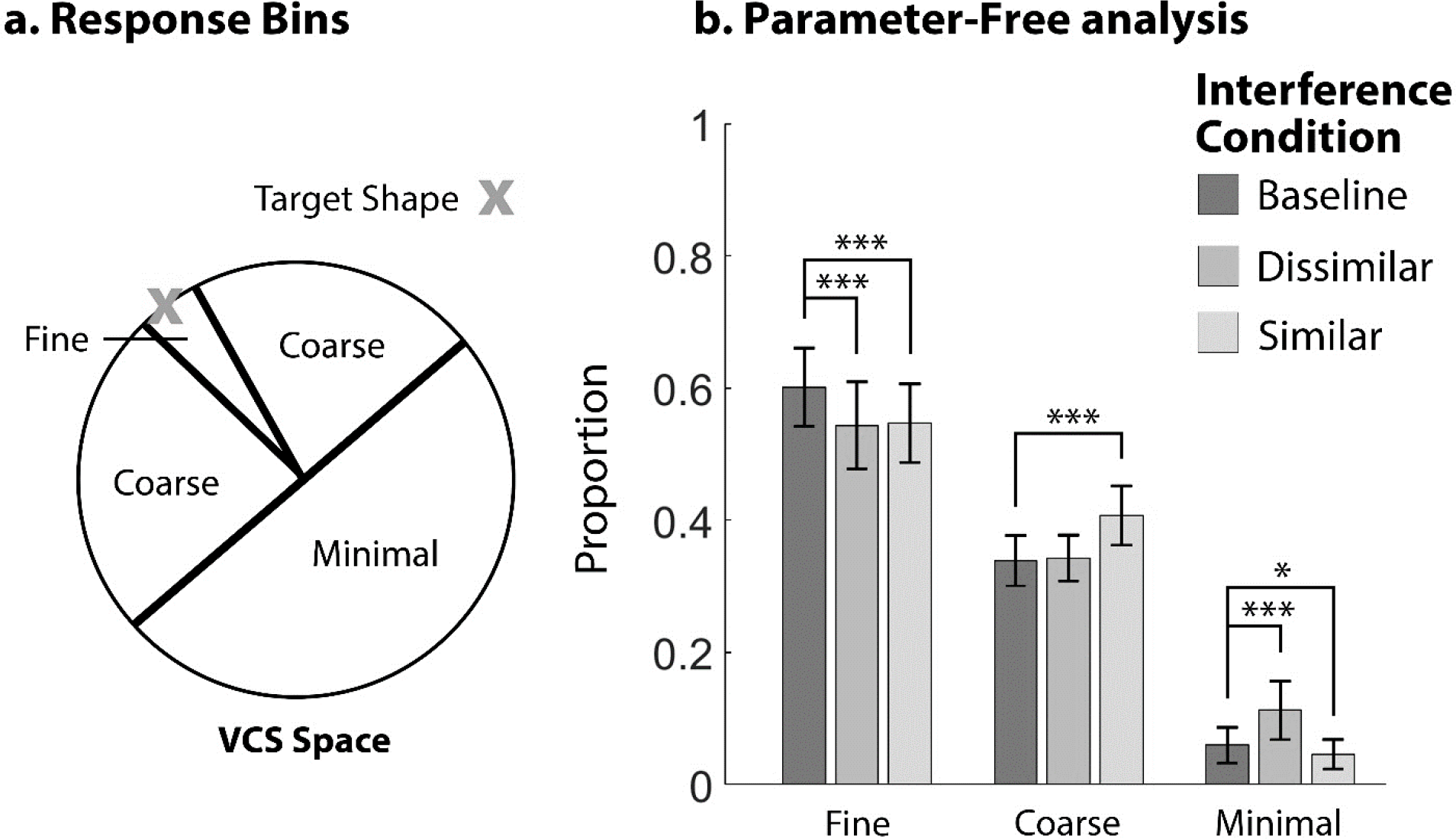
Parameter-free analysis. (a) Responses containing fine-grained, coarse-grained, and minimal target information were determined by binning regions of shape space. Fine-grained responses ranged from 14 degrees on either side of the target. Coarse-grained responses ranged from 15 – 89 degrees on either side of the target. Responses with minimal fine- or coarse-grained detail ranged from 91 degrees to 180 degrees on either side of the target. (b) The probability of reporting a fine-grained response was significantly lower following both dissimilar (distractors distal on circular space) and similar interference (distractors proximal on circular space) relative to a baseline condition consisting of scrambled lines. The probability of reporting a coarse-grained response was significantly higher in the similar relative to the baseline condition. The probability of reporting a response with minimal target information was significantly higher in the dissimilar relative to the baseline condition, and significantly lower in the similar relative to the baseline condition. Error bars reflect the 95% between-subject CI for the condition means. *** denotes *p* < 0.001, whereas * denotes *p* < 0.05.

The proportion of fine-grained responses was differentially influenced by interference condition (*F*_2,54_ = 11.99, *p* < 0.001, *partial R*^2^ = 0.31). The proportion of fine-grained responses was lower in both the dissimilar condition (*t*_54_ = 3.85, *p* < 0.001, *d* = 0.33) and the similar condition (*t*_54_ = 4.43, *p* < 0.001, *d* = 0.34) compared to the baseline condition. Thus, both dissimilar and similar interference disrupted fine-grained information.

The proportion of coarse-grained responses was differentially influenced by interference condition (*F*_2,54_ = 12.91, *p* < 0.001, *partial R*^2^ = 0.32). The proportion of coarse-grained responses was not significantly different between the dissimilar and baseline conditions (*t*_54_ = 0.30, *p* = 0.77). In contrast, the proportion of coarse-grained responses were significantly higher in the similar compared to the baseline condition (*t*_54_ = 4.77, *p* < 0.001, *d* = 0.58). Thus, similar interference increased the reliance on coarse-grained information.

The proportion of responses with minimal target-related information was differentially influenced by interference condition (*F*_2,54_ = 14.07, *p* < 0.001, *partial R*^2^ = 0.34). The proportion of responses containing minimal target-related information was significantly higher in the dissimilar compared to the baseline condition (*t*_54_ = 4.72, *p* < 0.001, *d* = 0.52). In contrast, the proportion of responses with minimal target-related information was significantly lower in the similar compared to the baseline condition (*t*_54_ = 2.42, *p* = 0.019, *d* = 0.20).

Critically, results from the parameter-free analyses replicated those from the mixture model analyses if we interpret retrieval success and precision along a continuum of fine- and coarse-grained information. One could consider retrieval success in terms of the relative proportion of responses with either fine- or coarse-grained target information, as opposed to responses with minimal information about the target. Likewise, one could consider precision to reflect the relative proportion of responses with fine-grained, as opposed to coarse-grained, information about the target. For example, after dissimilar interference, the decrease in retrieval success with no influence on precision (mixture model analyses) was consistent with the relative increase in responses containing minimal information about the target (parameter-free analyses). Likewise, after similar interference, the decrease in precision (mixture model analyses) was consistent with the relative increase in responses containing coarse-grained information about the target (parameter-free analyses). Finally, after both dissimilar and similar interference, the reduction in responses containing fine-grained information was met with either a relative increase in responses containing minimal information about the target (after dissimilar interference), or a relative increase in responses containing coarse-grained information about the target (after similar interference). In this manner, we interpret both mixture model parameters along a continuum of fine- and coarse-grained detail. We discuss the potential mechanisms underlying how the type of interference impacts the detail of representations in the Discussion (see the section *What Are the Mechanisms Underlying Forgetting?*).

## Discussion

Here, we provide converging evidence towards a generalizable set of principles governing how the type of interference impacts the detail of memories. Applying VCS space (Li et al., 2020) for the first time to a cognitive task, we explicitly quantified the visual similarity of distracting information and then measured how this impacted shape memory along a continuum of representational detail. We found that dissimilar distractors disrupted both fine- and coarse-grained information about the target, whereas similar distractors disrupted fine-grained information about the target but increased memory for coarse-grained information about the target. Critically, these results were consistent across multiple analyses: the mixture model fit to responses either scaled by angular distance or by perceived visual similarity to the target, and a parameter-free analysis that directly assessed fine- and coarse-grained information (*Figure 4*, 5, 6). Furthermore, these results were consistent with previous findings for color memory (Sun, Fidalgo, et al., 2017), confirming that the similarity of interference differentially and robustly impacts the representational detail of memory.

To characterize representational detail, we used two mixture model analyses (*Figure 4, 5*) and one parameter-free analysis (*Figure 6*). Whereas the mixture model is typically used to differentiate between retrieval success and the precision of the retrieved memory (*Figure 2*; Zhang & Luck, 2008), we have interpreted both parameters along a continuum of representational detail. Here, retrieval success was defined as the relative probability of remembering some fine- or coarse-grained information about the target, as opposed to a memory with minimal or no detail about the target. In a similar vein, precision was defined as the relative probability of remembering more fine-grained as opposed to coarse-grained information about the target. Recent debate has suggested that retrieval success and precision on the mixture model are not independent, with the distinction between these parameters resulting from an artefact of the non-linear relationship between angular distance and the magnitude of visual similarity on circular space (Schurgin et al., 2019). Here, we show that the overall conclusions obtained from the mixture model are replicated regardless whether responses were scaled by angular distance (*Figure 4*) or by the magnitude of visual similarity between two points on circular space (*Figure 5*). These results were furthermore consistent with a set of parameter-free analyses that did not fit the mixture model (*Figure 6*).

Although analyzing our shape data using multiple scaling metrics did not influence the overall conclusions, the *absolute* value of each parameter fit by the mixture model was impacted by the scaling metric. In other words, the numerical values of retrieval success and precision changed depending on whether responses were scaled by angular distance or by the magnitude of visual similarity. For example, whereas the precision parameter for the baseline condition was 0.065 when fit to errors scaled by angular distance (*Figure 4*), the precision parameter for the baseline condition was 0.036 when fit to errors scaled by the magnitude of visual similarity (*Figure 5*). Critically, the *relative* comparison between each interference condition with the baseline condition was not influenced by the scaling metric. In other words, errors from each experimental condition (baseline, dissimilar, and similar) were impacted by the approximation between angular distance and the magnitude of visual similarity in a consistent manner. Comparing each experimental condition against baseline therefore enabled us to examine the relative change to memory following interference, without making claims regarding the absolute amount of fine- or coarse-grained detail present in the target representation.

Regardless of the scaling metric, our results show that continuous measures of representational detail can be more informative than dichotomous yes/no metrics of accuracy and may be one way to reconcile the seemingly contradictory findings within the interference literature. More specifically, similarity may either help or hinder memory depending on the nature of the representations needed to perform a task. In tasks where coarse-grained information is sufficient to identify the target, similar distractors may serve as a coarse-grained reminder of the target item, thus benefitting memory (Jiang et al., 2016; Lin & Luck, 2009; Mate & Baques, 2009; Morey, 2018; Sanocki & Sulman, 2011). For example, researchers have shown that semantically similar distraction can help memory when coarse-grained category information about the target is cued (e.g., hearing the word “car part” helped memory for the target “tire” when in the presence of the distractor “termite”; Hanczakowski, Beaman, & Jones, 2017). However, in tasks where fine-grained information is necessary to identify the target, similar distractors may instead impair memory performance because the coarse-grained information can interfere with the fine-grained target memory (Baddeley, 1964; Baddeley & Dale, 1966; Barense et al., 2012; Cann, McRae, & Katz, 2011; Gordon, Hendrick, & Levine, 2002; Neely & LeCompte, 1999; Roediger & McDermott, 1995; Yeung et al., 2013; Watson & Lee, 2013; Wickelgren, 1965; Wickelgren, 1966; Wickens, 1970). For example, when distractors and targets were drawn from the same category, thus rendering coarse-grained categorical information no longer useful as a reminder about the target, semantically similar distraction caused memory impairment (e.g., hearing the word “car part” impaired memory for the target “tire” when in the presence of the distractor “wheel”; Hanczakowski, Beaman, & Jones, 2017).

Indeed, continuous measures of memory have driven a wealth of behavioral findings over the past decade (Ma et al., 2014). Simultaneously, there has been increasing interest in the neuroanatomical signatures of representational detail. Recent neuroimaging experiments suggest that different aspects of complex scenes can be represented concurrently, such as the fine-grained information about the color of objects embedded within scenes (Richter et al., 2016). Moreover, fine- and coarse-grained detail about complex scenes are represented within the medial temporal lobes and prefrontal cortex (Cooper & Ritchey, 2019; Kolarik, Baer, Shahlaie, Yonelinas, & Ekstrom, 2018; Nilakantan, Bridge, VanHaerents, & Voss, 2018; Stevenson et al., 2018; Yonelinas, 2013), perhaps modulated by differences in the neural dynamics associated with fine- and coarse-grained spatial information (Brunec, Moscovitch, & Barense, 2018; Bush, Barry, & Burgess, 2014; Collin, Millivojevic, & Doeller, 2015; Hasselmo et al., 2017; Poppenk, Evensmoen, Moscovitch, & Nadel, 2013). In one task, participants reconstructed the color, orientation, and spatial position associated with real-world objects studied on a 2D circle (Richter et al., 2016). Using a continuous retrieval task in conjunction with a mixture model, the experimenters differentiated between retrieval success and precision. Trial-wise activity within the hippocampus correlated with retrieval success (i.e., the relative proportion of responses with some fine- or coarse-grained information as opposed to minimal target-related information), whereas the angular gyrus correlated with the precision of memory (i.e., the relative proportion of responses with fine-grained as opposed to coarse-grained information about the target). These experiments have all made fruitful advances to the neuroanatomical understanding of memory, though researchers are often limited by the kinds of stimuli and tasks that can capture representational detail. Existing research typically use simple color and oriented line stimuli (Ma, Husain, & Bays, 2014) or capture the spatial distance between objects placed on a 2D circle (Berens, Richards, & Horner, 2020; Nilakantan, Bridge, VanHaerents, & Voss, 2018; Richter et al., 2016). Here, we provide empirical evidence that VCS space can be used not only to capture shape memory along a continuum of fine- and coarse-grained detail, but can also characterize the visual similarity of target and distractor shapes, thus offering experimenters a new tool for future work in this domain.

### What Are the Mechanisms Underlying Forgetting?

Throughout this manuscript, we have suggested that dissimilar distractors decreased both fine- and coarse-grained information, akin to “memory erasure”. We have furthermore suggested that similar distractors decreased memory for fine-grained information while increasing reliance on coarse-grained information, akin to “memory blurring”. Here, it is important to emphasize that “memory erasure” and “memory blurring” are metaphors for the complex processes that govern how interference impacts memory. Although there is clear evidence that distractor similarity differentially impacted representational detail (*Figure 4, 5, 6*), the mechanisms by which memory representations were altered by interference are much less clear. A full treatment of all the mechanisms underlying forgetting is well beyond the scope of this paper, but below we consider several possibilities, many of which are not mutually exclusive.

Following dissimilar interference, the reduction in retrieval success (mixture model analyses) driven by the relative increase in responses containing minimal target information (parameter-free analyses) could reflect not only the complete erasure of the target memory (Zhang & Luck, 2008; 2009), but also other mechanisms that can render the memory temporarily inaccessible (Tulving, 1974). One possibility is that the memory trace could not be accessed due to competition from distractors (Cowell, Bussey, & Saksida, 2006; Sadil & Cowell, 2017; Schurgin et al., 2019), resulting in errors made to competitors displayed throughout the experiment (e.g., also known as “swap”, “misbinding”, or “temporal binding” errors; Bays, Catalao, & Husain, 2009; Chun, 1997; Diana, Peterson, & Reder, 2004; Sadil & Cowell, 2017; Schurgin, Wixted, & Brady, 2019). Moreover, participants provide their response along a 2D circle during the test phase on a continuous retrieval task (*Figure 3*). Thus, not only do observers see distractors during the interference phase, observers search for the target during the test phase and are subsequently exposed to additional uncontrolled interference from shapes on the circular space which can render the memory inaccessible.

After similar interference, the reduction in precision (mixture model analyses) was driven by a relative increase in responses containing coarse-grained target information (parameter-free analysis), which we interpret as the “blurring” of the memory trace. One possible mechanism for this phenomenon is the direct distortion of the target memory by distracting information. It is well established that observers can retain the statistical regularities present in many different kinds of stimuli at a brief glance, such that individuals can recall the average color of an array, the average direction of pointed lines, or even the average emotional state of complex faces – all without fine-grained memory of individual items (Fischer et al., 2020; Rademaker, Bloem, De Weerd, & Sack, 2015; Whitney & Yamanashi Leib, 2018). Thus, each displayed distractor may be incrementally averaged with the target item during interference, leading to a bias in the memory for the target. Additionally, similar distractors may serve as a series of imprecise reminders which lead to disruptions in fine-grained memory about the target, with parallels to the phenomenon of retrieval-induced forgetting (Anderson, Bjork, & Bjork, 2000; Fukuda, Pall, Chen, Maxcey, 2020; Storm & Levy, 2012). Furthermore, rather than the memory of the target directly distorted by similar interference, a separate representation of the shared information between the distractors and targets may be created, which would also manifest as “blurring” (e.g., a categorical representation; Bae et al., 2015, Ester, Sprague, & Serences, 2020; Hardman, Vergauwe, & Ricker, 2017). Finally, as we described above for dissimilar interference, participants may make “swap” errors to similar distractors observed throughout the experiment. In this case, swaps to similar items would manifest as reductions in precision and an increase in responses to items bearing coarse-grained similarity to the target (Bays, Catalao, & Husain, 2009; Cowell, Bussey, & Saksida, 2006; Diana, Peterson, & Reder, 2004; Sadil & Cowell, 2017; Schurgin, Wixted, & Brady, 2018). Importantly, the behavioral manifestations of these mechanisms may be effectively indistinguishable on tasks such as ours. Perhaps even the underlying theories themselves share more similarities than differences, providing an intriguing direction of future work.

### Interpreting Mixture Model Parameters Along a Representational Continuum

What can the experimenter take away from the present debate regarding the interpretation of mixture models? Current debate in the short-term memory literature has revealed the difficulty in delineating the true absence of the memory trace (i.e., a complete lack of fine- and coarse-grained detail about the target) from the existence of a very weak memory trace (i.e., a representation with minimal detail about the target). Experimenters have suggested that there is no distinction between retrieval success and precision as captured by the mixture model, and instead proposed a single-parameter model that accounts for scaling differences between angular distance and the magnitude of visual similarity between points on circular space (Schurgin et al., 2019). Here, we reveal novel evidence that the conclusions from the mixture model are invariant to the scaling metric (*Figure 4, 5*), with the mixture model reliably characterizing the relative proportion of responses with minimal or no detail (i.e., *retrieval success*, captured through the uniform distribution) and the relative proportion of responses with fine-grained as opposed to coarse-grained information (i.e.,*precision*, captured through the von Mises distribution). Although we demonstrate the replicability of the mixture model across different scaling metrics, our interpretation of the mixture model parameters along a continuum of representational detail does not consider these parameters to be entirely independent. Instead, we show that the similarity of interference differentially impacts memory for fine- and coarse-grained information across both mixture model and parameter-free analyses (*Figure 4, 5, 6*). Future research should explore whether the current implementation of a one-parameter model based on signal detection theory can account for our data (Schurgin et al., 2019), as this model represents the entire distribution of errors as a single numerical value (e.g., “memory strength”) rather than as separate fine- and coarse-grained components that are free to vary. For example, a hypothetical finding of reduced “memory strength” after dissimilar compared to similar interference does not capture the full differences between fine-grained, coarse-grained, and minimal target-related information observed in our parameter-free analysis (*Figure 6*). In scenarios where fine-, coarse-grained, and minimal information are differentially impacted by experimental conditions, perhaps multiple parameters would be needed to successfully model this data.

Future investigations are needed to explore the relationship between the neural code, representational detail, and mixture model parameters. For example, how are neural representations lost during “memory erasure” after dissimilar interference or lost during “memory blurring” after similar interference? What are the neural codes supporting a fine- or coarse-grained representation for simple compared to more complex stimuli? Are these neural codes the same or different depending on the kinds of tasks used to assess memory? From further exploring these questions, perhaps we can uncover conditions in which certain models are better suited for understanding certain types of stimuli or certain types of tasks. For example, perhaps high-dimensional associative representations, such as real world objects and context, may be more usefully modelled in a discrete manner using a retrieval success parameter indicative of all-or-nothing forgetting (Berens, Richards, & Horner, 2020; Ekstrom & Yonelinas, 2020; Horner, Bisby, Bush, Lin, & Burgess, 2015; Joensen, Gaskell, & Horner, 2018). In contrast, low-dimensional representations such as single features (color, oriented lines, or shape) may be more usefully modelled in a continuous manner using a parameter which captures the representational detail of the memory trace (Ma et al., 2014; Schneegans & Bays, 2017; Schurgin et al., 2019).

Overall, we show that mixture model parameters reliably characterize the proportion of memories with some fine- or coarse-grained as opposed to minimal or no information (i.e., retrieval success) and memories with proportionally more fine-grained compared to coarse-grained information (i.e., precision), regardless of whether responses were scaled by angular distance or by the magnitude of visual similarity between items on a circular space. Interpreting mixture model parameters along a continuum of representational detail is furthermore consistent with parameter-free estimates which directly capture the magnitude of visual similarity. Future investigations may involve delineating whether retrieval success and precision as conceptualized on the mixture model are entirely independent, and whether our mixture model findings generalize to stimulus domains other than shape and color.

## Conclusion

The *Validated Circular Shape Space* (Li et al., 2020) was applied for the first time to quantify both the visual similarity of distracting information and the representational detail of shape memory. We found that the representational detail of memories was systematically and differentially altered by the similarity of distracting information. Dissimilar distractors disrupted both fine- and coarse-grained information about the target, akin to memory erasure. In contrast, similar distractors disrupted fine-grained but increased the reliance on coarse-grained information about the target, akin to memory blurring. As these effects have now been consistently observed across two stimulus domains of shape and colour, multiple scaling metrics, as well as multiple mixture model and parameter-free analyses, we suggest these results provide a general set of principles governing how the type of interference impacts forgetting.

## Acknowledgements

We thank Celia Fidalgo for her invaluable contributions to the development of this study and its interpretation, Timothy Brady, Stephen Rhodes, Sol Z. Sun, and Bryan Hong for helpful discussions, as well as Elizabeth Page-Gould for statistical assistance. AYL is supported by an Alexander Graham Bell Canada Graduate Scholarship-Doctoral from the Natural Sciences and Engineering Research Council of Canada (NSERC CGS-D). This work is supported by a Scholar Award from the James S McDonnell Foundation, an Early Researcher Award from the Ontario Government, an NSERC Discovery grant, and a Canada Research Chair to MDB. This work is also supported by an NSERC Discovery grant to ACHL.

